# Obesity alters mobility and adult neurogenesis, but not hippocampal dependent learning in ob/ob mice

**DOI:** 10.1101/537720

**Authors:** Alexander Bracke, Grazyna Domanska, Katharina Bracke, Steffen Harzsch, Jens van den Brandt, Barbara Bröker, Oliver von Bohlen und Halbach

**Affiliations:** Institute of Anatomy and Cell Biology, University Medicine Greifswald, Friedrich Loeffler Str. 23c, D-17489 Greifswald, Germany; Institute of Immunology and Transfusion Medicine, Department of Immunology, University Medicine Greifswald, Ferdinand-Sauerbruch-Straße / DZ 7, D-17475 Greifswald, Germany; Zoological Institute and Museum, Department of Cytology and Evolutionary Biology, University Greifswald, Soldmannstrasse 23, D-17498 Greifswald, Germany; Central Service and Research Unit for Laboratory Animals (ZSFV), University Medicine Greifswald, Walther-Rathenaustr. 49a, D-17489 Greifswald, Germany

**Keywords:** leptin, obesity, cognition, Morris water maze, hippocampus, ob/ob mice

## Abstract

**Background:** Obesity has become a severe problem among the world’s population with clearly increasing prevalence over the last decades. Because obesity is associated with several comorbidities (e.g. hypertension or cancer) it constitutes an increasing burden for the health care system. Correlations between obesity and cognition have been studied in humans with ambivalent results. Here, we studied the effects of obesity on hippocampus dependent learning and memory and cell morphology in a mouse model of obesity.

**Methods:** The body mass of male and female Lep^+/+^(wt) and Lep^ob/ob^(ob/ob) animals with access to food and water ad libitum was measured between postnatal day 60-200 and animals with clear adiposity (4-6 months) were further analyzed. Adult hippocampal neurogenesis in the dentate gyrus was examined using phosphohistone H3 as a marker for proliferation, doublecortin as a marker for differentiation and caspase3 as a marker for apoptosis. Moreover, the density of dendritic spines on apical and basal dendrites of pyramidal neurons of the cornu ammonis 1 (CA1) were analyzed using Golgi impregnation. In addition, mice were subjected to the open field and Morris water maze test in order to analyze locomotor activity and spatial learning.

**Results:** The body weight of ob/ob mice nearly doubled during the first 120 postnatal days. Adult hippocampal neurogenesis was reduced in ob/ob mice due to reduced cell proliferation. Dendritic spine densities in the hippocampal area CA1 were not altered in ob/ob mice. Four to six months old ob/ob mice showed reduced locomotor activity in the open field test but similar performance in the Morris water maze compared to control mice.

**Conclusion:** Our data show that alterations in adult neurogenesis in leptin-deficient mice are not associated with an impairment in spatial learning abilities. Moreover, ob/ob mice are inconspicuous in the Morris water maze and do not display altered spine densities in the hippocampus, suggesting that obesity does not have a severe impact upon hippocampal neuronal plasticity and spatial learning.

## Introduction

According to estimates of the World Health Organization (WHO) from the year 2016, 1.9 Billion of the world’s population (aged 18 years and over) could be classed as overweight or obese based on their body mass index (BMI) (1). The BMI is the ratio between body weight and squared body height (BMI = weight (kg)/(height (m))^2^). For adults a BMI considered normal is between 18.5 kg/m^2^ to 24.9 kg/m^2^. A BMI between 25 kg/m^2^ and 29.9 kg/m^2^ is considered overweight. With a BMI greater or equal to 30 kg/m^2^, adults are classed as obese (2).

Obesity is a major health issue in many countries of the world, since obesity contributes to premature deaths by increasing the risk of cardiovascular disease, diabetes mellitus type II and even some forms of cancer (3–5). According to the WHO today the majority of the world’s population lives in countries reporting more deaths caused by obesity than malnutrition (1). A possible relationship between obesity and cognition in humans has been studied with quite controversial results. Some studies have linked deficits in general cognitive performance (6), psychomotor performance and speed (7, 8), visual and verbal memory (8, 9), as well as verbal fluency (9) with obesity. However, other analyses failed to find such correlations (9–12). Nevertheless, some studies indicate obesity resulting in motor deficits (13, 14) and deficits regarding self-restraint (15, 16). Seeyave et al. Have demonstrated that eleven year old children were more likely to be obese, when diagnosed with impaired self-restraint at the age of four (17). According to this study, obesity could be, at least in some cases, be interpreted as a symptom of deficits in self-restraint. Furthermore, investigating the relationship of academic success and obesity also yielded controversial data. While some studies found a negative impact of obesity on academic success (18, 19), Barrigas et al. reported that such a correlation does not exist when the social and economic background of the participants were taken into account in the data analysis (20).

Interpreting and comparing different studies in humans is highly challenging, because most studies used different methodological approaches and also the parameters the participants are controlled for, such as age, sex, comorbidities (e.g. diabetes mellitus, hypertonia) or educational level, are not standardized. These aspects were recently discusses in the review by Christina Prickett (21). Henceforth, an unambiguous causality of obesity and cognition today is hard to prove, although some analyses seem to foster this interpretation (for more details see (21, 22). In general, studies involving humans are hampered by the fact that usually a genetically very heterogeneous population, composed of individuals with highly variable living conditions, is analyzed. To overcome this set-back, in the current study we used an established animal model of obesity to analyze the impact of obesity on brain morphology and function. To study the effects of obesity, leptin-deficient (Lep^ob/ob^) mice or leptin-receptor-deficient (Lep^db/db^) mice are established models. Both mouse lines are characterized by a strong weight gain within the first postnatal months. As they grow up, the leptin receptor deficient mice (db/db), develop a permanent form of diabetes mellitus, whereas the ob/ob mice do not (23, 24). Thus, in contrast to the diabetic db/db mice the blood glucose concentrations were not increased in the ob/ob mice (25). In order to avoid an impact of diabetes on brain morphology and / or function, we used the Lep^ob/ob^ mouse strain (ob/ob) to analyze effects of obesity on neuronal plasticity and hippocampal learning and memory. We focussed on adult hippocampal neurogenesis, dendritic spine densities in hippocampal area CA1 and the behavior of the obese mice in the open field and Morris water maze.

## Materials and Methods

### Animals

Ob/ob mice as well as wild type littermates (+/+) were generated by crossing of heterozygous B6.V-Lep^ob^/J (+/ob) animals, since homozygous female mice are infertile and males show reduced fertility. Animals were kept in a 12 hour day-night cycle with food and water access ad libitum. Leptin-deficient (ob/ob) and littermate wild type control (wt) mice of both sexes were used in all subsequent experiments.

In the experiments ob/ob mice and littermate wild type controls of both sexes were analyzed. For body weight analysis mice were 60-200 days old, all subsequent experiments were conducted with mice at an age of 4-6 months.

All experiments were performed according to institutional and/or national guidelines for animal care of the “Landesamtes für Landwirtschaft, Lebensmittelsicherheit und Fischerei Mecklenburg-Vorpommern (LALLF M-V)” (7221.3-1-016/16.)

### Body weight analysis

To determine the development of body weight 48 control and 48 ob/ob animals (from postnatal day 60 up to postnatal day 200) were weighed once a week using a Beurer KS 36 scale (Beurer, Germany).

### Microvolumetry

Brain volume was determined by using microvolumetry (µ-VM) as described previously (26). In brief, a 5 ml syringe was used as sample container with a 1 ml syringe attached as measurement device. 3 ml of 4 % paraformaldehyde (PFA) were used as fluid in the sample container in which the mouse brain was placed. Then the initial level of 3 ml was restored using the measurement device in which the displaced volume could be measured. Measurements were repeated three times for each brain (n = 19 per group) and averaged.

### Immunhistochemical analysis

Animals were euthanized and transcardially perfused with phosphate buffered saline (PBS) and 4 % PFA. The brains were explanted and stored in 4 % PFA at 4°C for four days. Thirty µm coronal sections were made using a vibration blade microtome (Type VT 1000 S, Leica, Germany) Sections were mounted on superfrost slides (R. Langenbrinck GmbH, Germany) and dried over night at 37°C. Rehydration and antigen retrieval was achieved via microwave treatment(Samsung Electronics GmbH, Germany; 20 min, 800 W).

For Phosphohistone H3 (PH3) staining sections were rinsed with distilled water (A. dest.) and PBS and unspecific binding sides were blocked using 5 % normal goat serum (NGS) + 5 % bovine serum albumine (BSA) + 0.1 % Triton X-100 in PBS for 60 min at room temperature (RT). After washing with PBS, primary rabbit anti-phosphohistone H3 antibody (Santa Cruz, USA) diluted 1:100 in 1 % NGS + 0.1 % Triton X-100 in PBS was applied and incubated for 120 min at 4°C. After washing with PBS, the secondary antibody (Cy3-conjugated goat anti-rabbit-IgG (DIANOVA, Germany) diluted 1:2.000 in 1 % NGS + 0.1 % Triton X-100 in PBS) was applied and specimen incubated for 60 min at RT. Afterwards, all sections were rinsed in PBS and counterstained with 4’6-diamidino-2-phenylindole dihydrochloride (DAPI, 1:10.000), then washed with A. dest. and embedded in Mowiol (Merck KGaA, Germany).

For doublecortin (DCX) staining, sections were permeabilized with 0.4 % Triton X-100 in PBS for 30 min. After washing with PBS, sections were incubated in blocking solution (3 % horse serum + 0.3 % Triton X-100 in PBS) for 60 min at RT. After rinsing with PBS, sections were incubated in a solution containing the primary goat anti-doublecortin antibody (1:100 (Santa Cruz Biotechnology, Germany) in 3 % horse serum + 0.1 % Triton X-100 in PBS), and incubated for 3 days at 4°C. Rinsing with PBS was followed by incubation in a solution containing biotin-conjugated horse anti-goat IgG (1:200 (Vector Laboratories, USA) in 3 % horse serum + 0.1 % Triton X-100 in PBS) for 2 hours at RT. After washing in PBS, the secondary antibody was applied to the sections (Cy3-conjugated Streptavidin (DIANOVA, Germany), 1:2.000 in PBS). Incubation time was set to 2 hours at RT. All sections were rinsed in PBS and counterstained with DAPI, 1:10.000 in A. dest., washed with A. dest and embedded in Mowiol (Merck KGaA, Germany).

For Caspase3 (Casp3) staining, sections were rinsed with A. dest. and PBS and blocking solution was applied (3 % NGS + 0.1 % Triton X-100 in PBS). Sections were incubated for 60 min at RT. After washing with PBS, the solution containing the primary rabbit anti-active Caspase 3 antibody (Merck Millipore, Germany) was applied 1:100 in blocking serum (3 % NGS + 0.1 % Triton X-100 in PBS) and incubated at 4°C over night. Rinsing the sections with PBS was followed by the application of the secondary antibody containing solution (Cy3-conjugated goat anti-rabbit-IgG (DIANOVA, Germany) 1:400 in blocking serum) and incubated at RT for 90 min. Afterwards all sections were rinsed in PBS and counterstained with DAPI (1:10.000 in A. dest.), washed with A. dest and embedded in Mowiol (Merck KGaA, Germany).

We analyzed immunostained cells within the dentate gyrus (DG) using an Axioplan 2 imaging microscope (Zeiss, Germany) with a 400x magnification (40x objective + 10x ocular). All DCX (wt n = 13; ob/ob n = 9), PH3 (wt n = 9; ob/ob n = 10) and Casp3 (wt n = 12; ob/ob n = 8) positive cells were counted during microscopic analysis through all sections and averaged. The number of positive cells in wild type animals was normalized to 100 %.

## Analysis of dendritic spines

Animals were sacrificed (wt n = 6; ob/ob n = 6) and transcardially perfused with PBS and 4 % PFA (in PBS, pH 7.2). The brain tissue was stored in 4 % PFA at 4°C for at least three days for post-fixation. Then, we conducted a Golgi silver impregnation using the FD Rapid GolgiStain™ Kit (FD Neuro Technologies, USA). Coronal brain sections of 120 µm were made using a vibration blade microtome (Type VT 1000 S, Leica, Germany). Sections were mounted on gelatinized glass slides, dehydrated and exposed to Xylol. The mounted sections were coverslipped with Merckoglas® (Merck Millipore, Germany).

We analyzed dendrites of hippocampal CA1 pyramidal cells and focused on secondary or tertiary dendrites, as described previously (27). Only one segment per dendritic branch was chosen and approximately 20 apical and 20 basal dendrites were analyzed. The mean number of dendritic spines corresponds to a dendritic length of one micrometer. Using NeuroLucida 9.12 software (MBF Bioscience, USA) to control the x-y-z axis of the microscope (Axioscope Imaging, Zeiss, Germany) and a digital camera attached to it (AxioCam Hrc; Zeiss, Germany), we were able to reconstruct the dendrites 3-dimensionally using a 100x oil objective (NA: 1.4; oil immersion), as described previously (27). The reconstructed dendrites were then analyzed using the software NeuroExplorer (MBF Bioscience, USA).

Statistical analysis was based on the animal numbers (n = 6 for both groups) and not on the numbers of reconstructed dendritic spines (we analyzed about 4827 ± 475.18 dendritic spines of apical and about 4653 ± 110.31 dendritic spines of basal dendrites of pyramidal cells in the CA1 region of the hippocampus per group).

## Behavioral analysis

For behavioral analyses mice were subjected to two different tests.

(a) Open field (OF): A quadratic 45 x 45 cm test arena (Panlab, Spain) was used for the open field test, with illumination set to 25 lux. To start a trial, the animal (wt n = 29; ob/ob n = 32) was placed in the center of the area for 7 minutes to explore the environment. The tracking started automatically and movements were recorded via webcam (Logitech C300, Switzerland). Parameters characterizing open-field behavior (total distance covered (cm) and velocity (cm/s)) were analyzed from recorded sessions using SmartJunior 1.0.0.7 (Panlab, Spain). Between trials the arena was cleaned with 70 % ethanol.
(b) Morris water maze (MWM): Animals (wt n = 26; ob/ob n = 26) were trained to localize a circular, hidden platform (Ø 11 cm, acrylic glass; 2 cm underneath the water surface) within a water-filled circular pool (Ø 140 cm). Therefore, animals could only use visual cues of simple geometric figures attached to the surrounding walls of the room. The water (renewed daily) was rendered opaque with milk powder (550 g in 740 l water; Basu Mineralfutter GmbH, Deutschland) and kept at 24°C ± 1 °C. Illumination was set to 25 lux. Animals were trained in 20 trials, with 4 trials on 5 consecutive days (inter-trial interval 30 min) during which the position of the platform was kept unchanged (acquisition phase). The starting position for each animal was changed between trials. During the acquisition phase, mice were allowed to explore the pool for a maximum of 120 s or until finding the platform. If they failed, they were guided onto the platform. On the sixth day (probe trial) the platform was removed from the water maze. Mice were allowed to explore the water maze for 120 s.

After one day of resting, animals had to perform in the spatial reversal task (28–30). The platform position was moved to the opposite side of the pool to test the ability of the animals to adapt to the altered test situation. The training phase and probe trial were then performed as described above (4 training sessions on 5 consecutive days, probe trial on the 6th day).

With a webcam (Logitech C905, Switzerland) centered above the pool, swimming tracks of each individual were recorded and analyzed using Smart 3.0 (Panlab, Spain). The following parameters were recorded by the software during the acquisition phase: swimming speed (cm/s) and time to target platform (s); during the probe trial the software additionally recorded the parameters: platform position crossings and time in platform quadrant (s). The tracking data also allowed us to calculate an additional parameter, which was independent from the generally mobility-impaired velocity of the animals. The parameter “swimming direction” (%) was defined as the swimming accuracy, by measuring the speed-independent movement towards the platform position.

## Statistics

GraphPad Prism 5.0 (GraphPad Software Inc., USA) was used for statistical analysis of all data. We conducted unpaired t tests on all data with a level of significance set to p ≤ 0.05. Data in the figures were expressed as mean ± standard error of the mean (SEM). Significant changes are labeled as * p ≤ 0.05, ** p ≤ 0.01, *** p ≤ 0.001.

## Results

### Weight nearly doubles in ob/ob mice during the first 120 postnatal days

Leptin-deficient mice show an obese phenotype due to the defective regulation of food intake. However, this phenotype continuously develops over time. We therefore monitored up to 48 individuals of each genotype from postnatal day 60 until postnatal day 200. We were especially interested in determining the age at which obese mice display an average body weight twice as high as wild type control animals. We observed that at postnatal day 120, the weight of the obese mice was doubled (55.1 g; Fig. 1) as compared to the control mice (26.4 g).

**Fig. 1.**
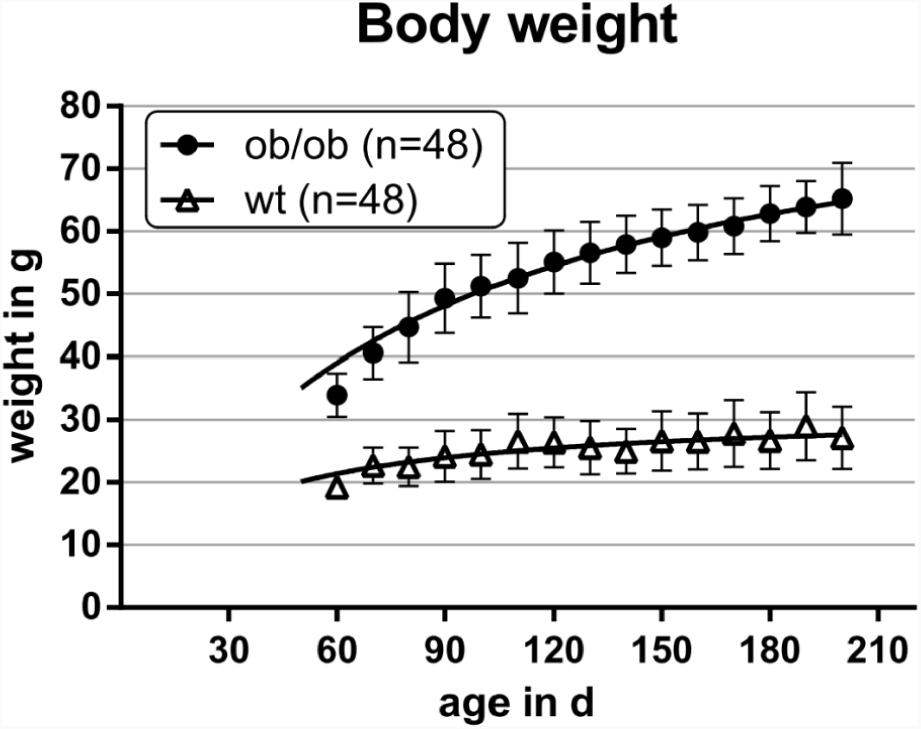
Body weight of wild type (wt) and ob/ob mice was measured from postnatal day 60 until postnatal day 200. Leptin-deficient mice show a twofold body weight in comparison to control animals with about 120 days of age. Data from 10 consecutive days were pooled (n = 48 per group).

### Brain volume is reduced in ob/ob mice

For analyzing the whole brain volume of ob/ob mice and wild type animals (n = 19 per group) we used microvolumetry. Adult ob/ob mice display a significantly reduced brain volume of about 10.7 % as compared to age-matched wild type controls (Fig. 2).

**Fig. 2.**
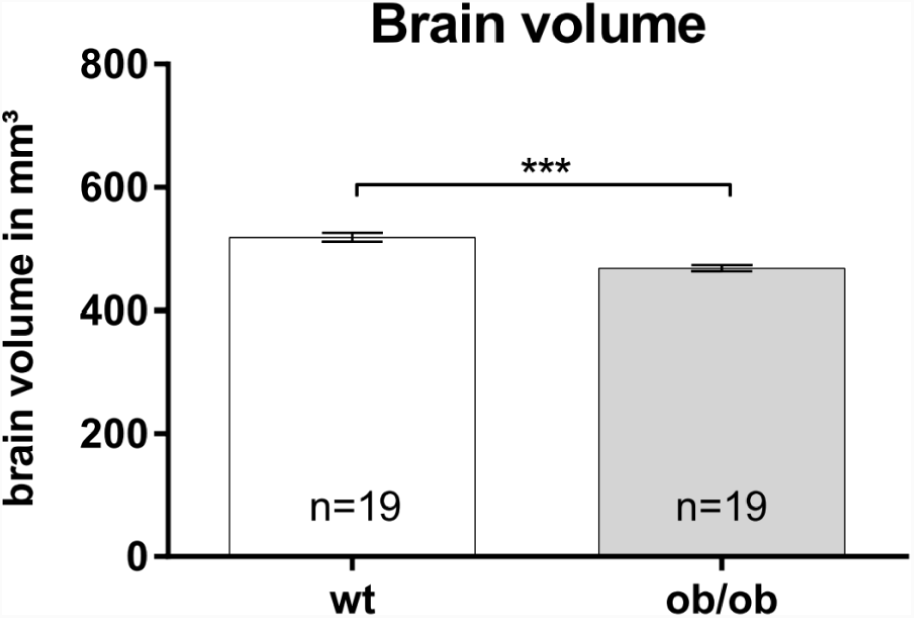
Whole brain volumes were analyzed using microvolumetry. Brains of leptin-deficient mice (ob/ob) are significantly smaller as compared to wild type (wt) controls (n = 19 per group).

### Adult hippocampal neurogenesis is altered in ob/ob mice

Since neurogenesis can be classified into different phases such as, e.g., stages of proliferation, differentiation, migration, maturation and synaptic integration, we used immunofluorescence staining of marker proteins that are being expressed in distinct phases of the development of new born neurons (31).

Antibodies directed against PH3 allow to label proliferating cells in m-phase. The number of PH3 positive cells (Fig. 3A) in the DG is significantly reduced by ∼ 18 % in ob/ob mice (n = 10) as compared to control littermates (n = 9). PH3 is specific for mitotic cells, but is not specific for newly formed neuronal cells (mitotic as well as postmitotic). In order to analyze this cell population we used the marker DCX and determined the number of DCX-positive cells in the DG. The analysis revealed that ob/ob mice (n = 9) show significantly less (∼ 32 %) DCX positive cells in the DG as compared to controls (n = 12; Fig. 3B). Because during adult neurogenesis, proliferation and differentiation is accompanied by apoptosis, we next analyzed whether there are differences in apoptosis in the DG by using antibodies directed against active (cleaved) casp3. The analysis revealed no significant differences between wt (n = 12) and ob/ob mice (n = 8; Fig. 3C).

**Fig. 3.**
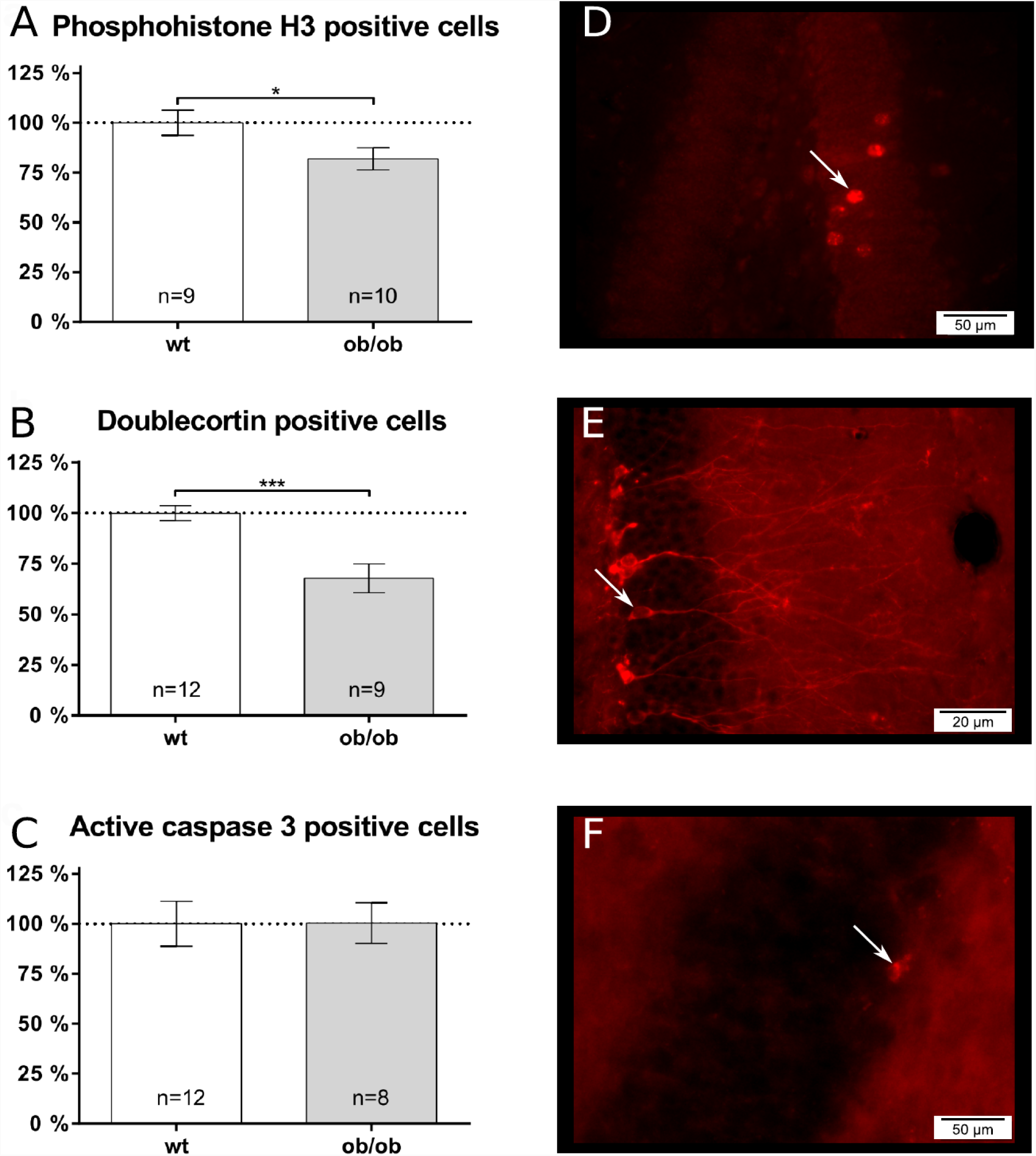
Adult neurogenesis, proliferation and apoptosis were analyzed using immunofluorescence staining within the dentate gyrus (DG) of the hippocampus. Cell numbers of wild type (wt) animals were set to 100 %. **A** The number of proliferating cells was visualized using phosphohistone H3 (PH3). Leptin-deficient (ob/ob) mice showed a reduced amount of PH3-positive proliferating cells. **B** As a maker for immature neuronal cells, doublecortin (DCX) was used. Ob/ob mice show a reduced number of DCX positive cells in the DG compared to wild type controls. **C** As an apoptosis marker caspase3 (Casp3) was used. The number of Casp3 positive cells did not differ between the two groups of mice. Fluorescence images were taken using a Olympus BX63 microscope with 25,2x magnification (40x objective, 0,63x camera) in the DG of wt mice for **D** PH3 positive, **E** DCX positive and **F** Casp3 positive cells.

### Dendritic spine densities in area CA1 are not altered in ob/ob mice

We analyzed the density of dendritic spines on apical and basal dendrites of pyramidal neurons in the hippocampal area CA1. Although elevated spine densities were noted on apical dendrites (wt: 1.193 ± 0.036 versus ob: 1.367 ± 0.074; Fig. 4A) and basal dendrites (wt: 1.137 ± 0.027 versus ob: 1.206 ± 0.064; Fig. 4B) of ob/ob mice, these differences did not reach statistical significance.

**Fig. 4.**
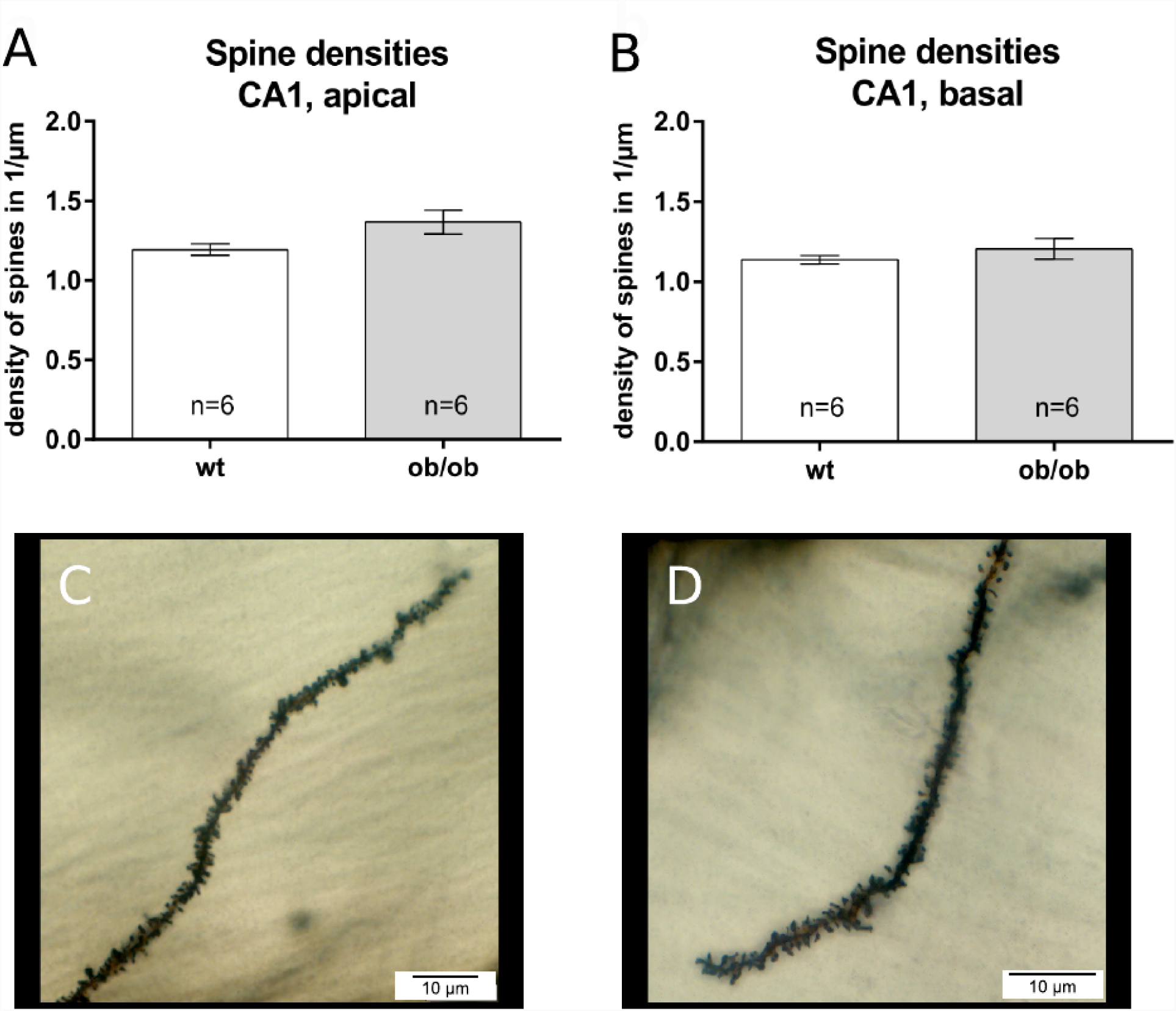
Analyzing the density of dendritic spines revealed no significant differences but showed a non-significant trend towards increased spine densities in Leptin-deficient (ob/ob) animals of **A** apical and **B** basal dendrites of pyramidal cells in the CA1 region of the hippocampus. Microscopic images show apical dendrites of **C** wild type (wt) and **D** obese mice of CA1 pyramidal cells. Images were taken using a Olympus BX63 microscope with 63x magnification (100x objective, 0.63x camera).

### Locomotor activity is reduced in ob/ob mice

The open field test (OF) is a common and standardized test to analyze the general locomotor activity of mice. The severe obesity should result in a decreased locomotor activity. Therefore, we wanted to measure how incisive this limitation of locomotor activity was in four to six months old ob/ob mice and their wt controls. In the OF general locomotor activity was determined by analyzing the distance the animals traveled within 7 minutes. Wild type animals (n = 29) traveled an average of 3817 ± 153.3 cm, whereas ob/ob mice (n = 32), within the same time, show a significant difference by traveling about 43.2 % less (1648 ± 116 cm; Fig. 5A). In addition, the ob/ob mice display a significant reduced average velocity over time of about 43.5 % (wt: 9.023 ± 0.3613 cm/s versus ob/ob: 3.925 ± 0.2716 cm/s; Fig. 5B).

**Fig. 5.**
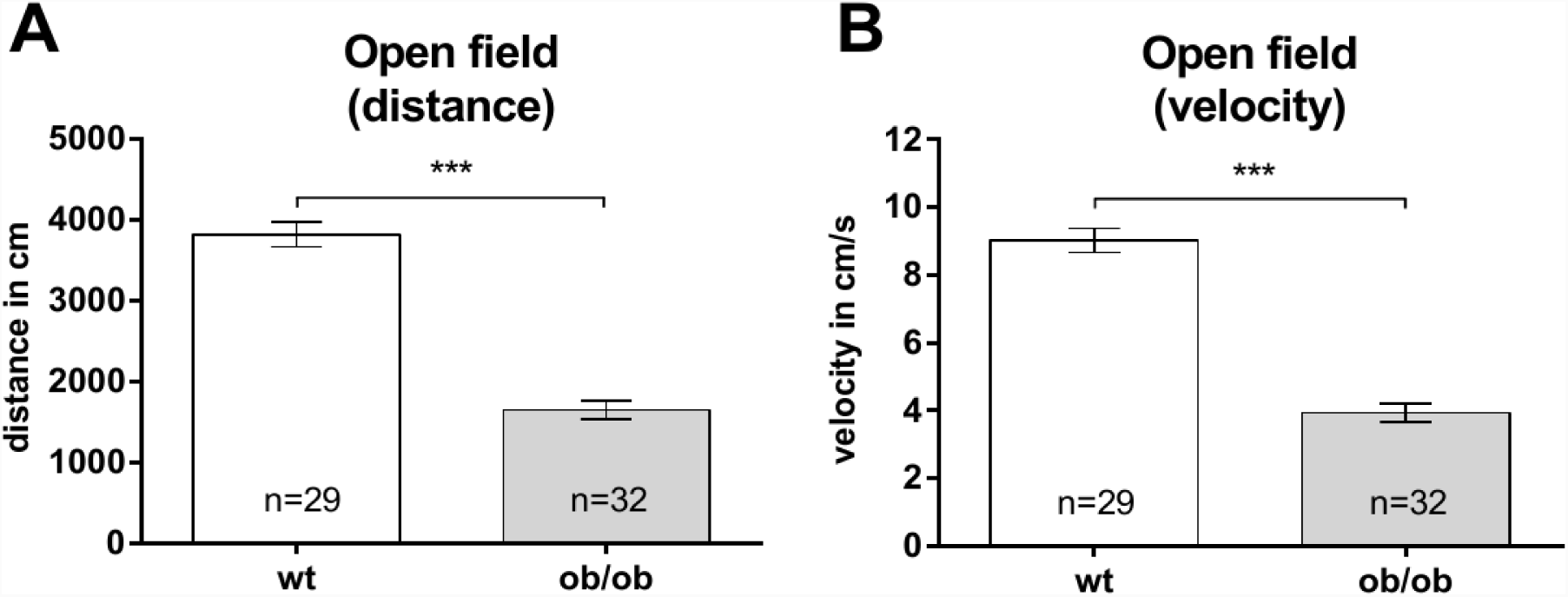
To analyze the locomotor activity of Leptin-deficient (ob/ob) mice, the open field test was used. **A** Ob/ob mice covered significantly shorter distances with **B** significantly reduced velocity compared to wild type (wt) mice.

### Ob/ob and wt mice show comparable performances in the Morris water maze test

To analyze the impact of obesity on learning abilities and memory function, we conducted the Morris water maze test in which the animals are being trained to locate a hidden platform within a water basin.

Both, ob/ob mice and their wt controls were able to learn the task during the training phase. In accordance to our data generated in the open field test, the ob/ob mice showed a significantly reduced swimming speed (Fig. 6A). However, during the first four days the mean time to the target platform was not significantly altered between both groups (Fig. 6B). Keeping in mind the reduced mobility of ob/ob mice, as shown by the open field data as well as demonstrated by activity measures by others (32), we calculated an additional parameter, which was independent from the mobility-impaired speed of the ob/ob animals. Analyzing the parameter “swimming direction” revealed that the performance in localizing the hidden platform of ob/ob mice was comparable to lean wt controls (Fig. 6C). After 5 consecutive training days the animals performed in a probe trial. In this probe trial the platform was removed from the water maze. Then we analyzed the latency to find the former platform position (wt: 13.78 ± 3.251 s versus ob/ob: 19.28 ± 3.428 s; Fig. 7A), as well as the time animals spent in the quadrant, where the platform used to be (wt: 38.8 ± 1.85 % versus ob/ob: 37.71 ± 1.871 %; Fig. 7B). Both parameters showed no significant differences between groups.

**Fig. 6.**
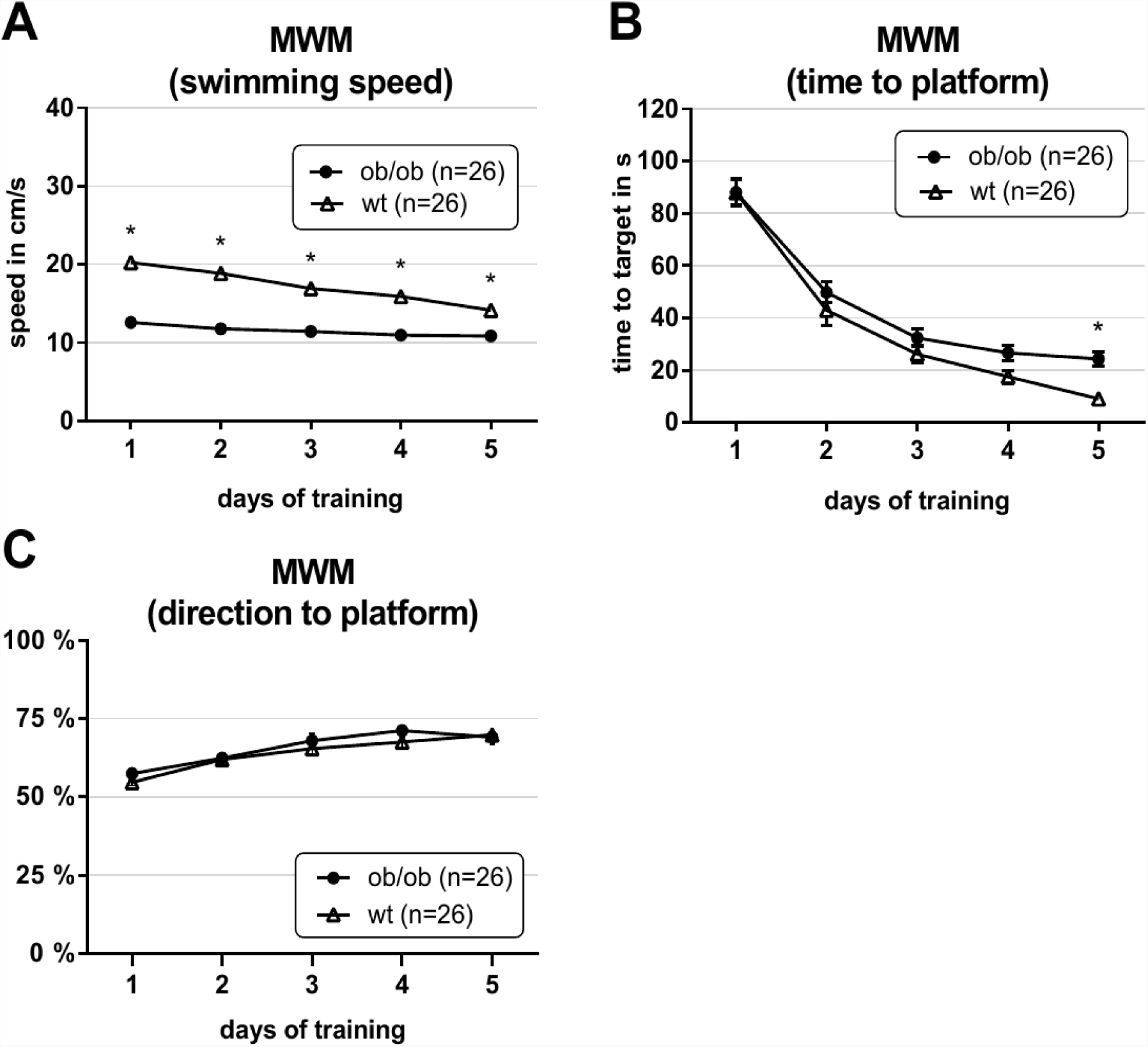
Leptin-deficient mice (ob/ob) show comparable results in localizing the hidden platform in comparison to age-matched wild type (wt) control mice. **A** Ob/ob mice displayed a significantly reduced swimming speed. **B** Concerning the time to reach the platform, ob/ob mice show overall comparable results to their controls. Only on day 5 mice exhibited a significant difference. **C** The swimming direction also showed comparable swimming accuracy towards the platform position in comparison to wt controls.

**Fig. 7.**
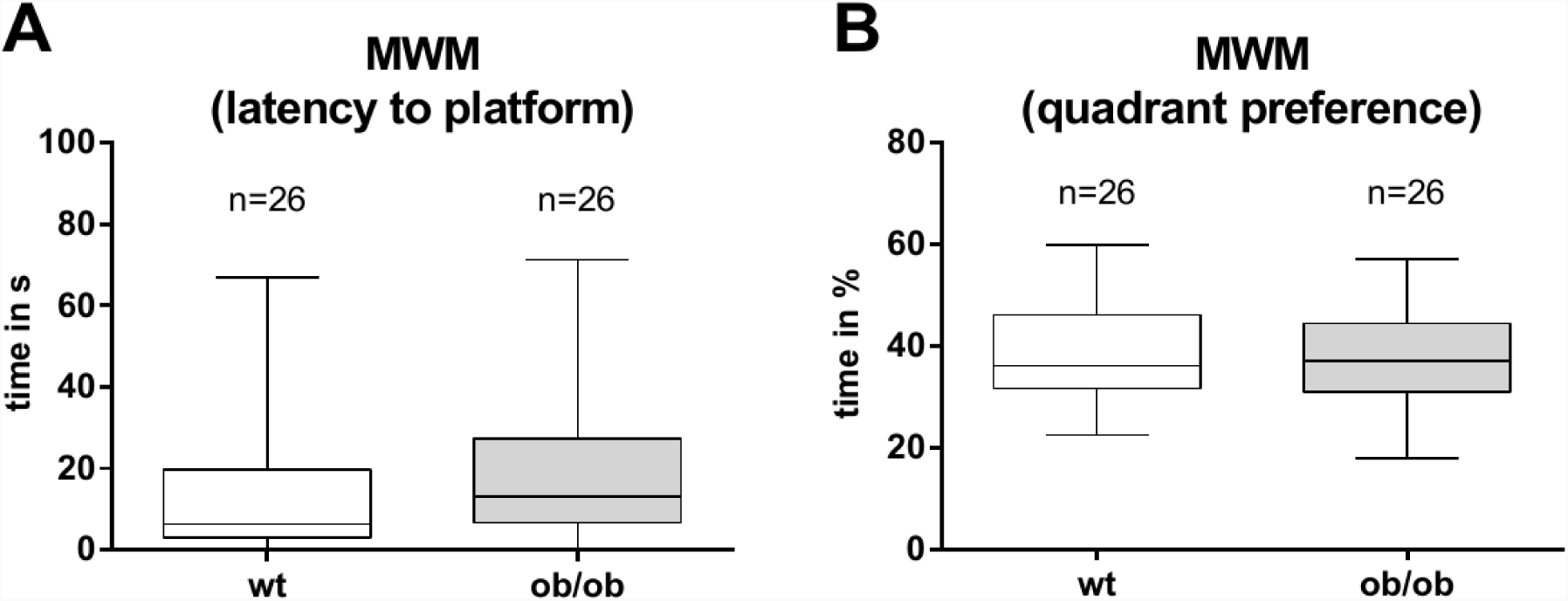
In the probe trial the Leptin-deficient (ob/ob) mice showed no significant differences regarding **A** the time to reach the platform position or **B** the time spent in the platform quadrant. Data are represented as boxplot with median line and whiskers for lowest and highest values.

After the first week the animals had to perform in the spatial reversal task (28, 29). The idea of this test is to slightly vary the already known behavioral task by placing the platform to a position of 180 degrees on the opposite side in the water maze. When testing the animals again it is possible to evaluate their ability to adapt to the changed situation. The spatial reversal also consists of a 5 day training and probe trial on the 6th day. Accordingly to the first week data the time to reach the platform was not significantly altered between both genotypes (Fig. 8A). The swimming speed was again significantly different in the second training phase (Fig. 8B). Surprisingly, ob/ob mice showed a more accurate swimming direction in the spatial reversal task (Fig. 8C).

**Fig. 8.**
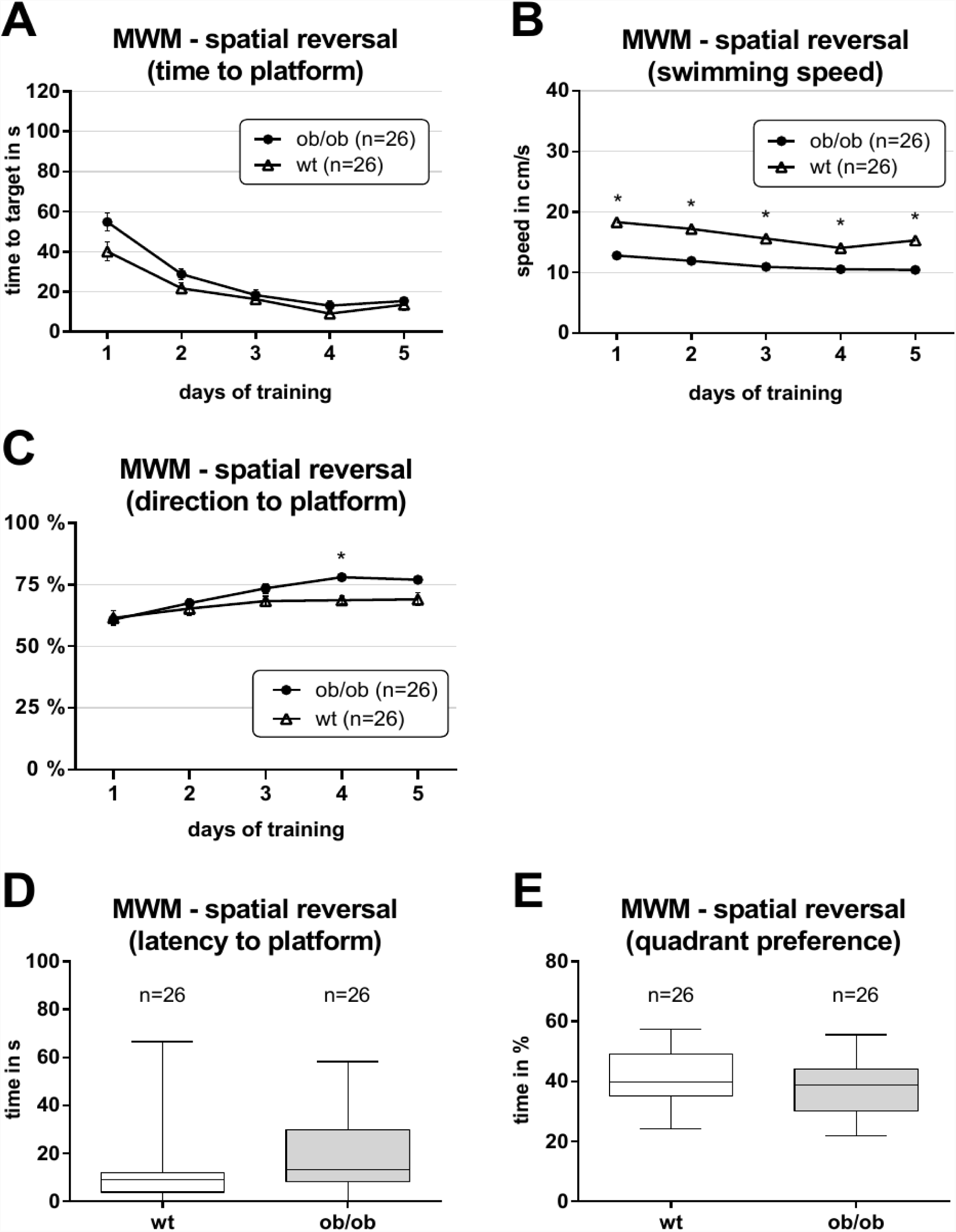
In the spatial reversal experiment, the platform is positioned on the opposite side of the maze. **A** Leptin-deficient (ob/ob) and wild type (wt) mice showed comparable latencies to reach the platform. **B** Ob/ob mice continued to show reduced swimming speed in the spatial reversal task. **C** Ob/ob mice were able to reach higher accuracy concerning the swimming direction towards the platform position. The data acquired for the probe trial did not differ for **D** the latency to reach the platform or **E** the preference of the platform quadrant. Data in **D** and **E** are represented as boxplot with median line and whiskers for lowest and highest values.

During the probe trial of the spatial reversal Leptin-deficient animals did not show significantly different performance to wild type controls concerning their latency to platform (Fig. 8D) or platform quadrant preference (Fig. 8E).

## Discussion

Several genetic mouse models of obesity are available (e.g. ob/ob mice, db/db mice) as well as diet-induced approaches (23). We have chosen a mouse model of obesity, based on leptin-deficiency (ob/ob), since at 4 months of age the ob/ob mice weight approximately twice as much as lean littermate control animals (see Fig. 1). In the present study, we used 4 to 6 month old mice to investigate the possible effects of obesity on neuronal plasticity, learning and memory. A related mouse model are the obese and diabetic leptin-receptor deficient mice (db/db) (23). In contrast to the ob/ob mice, young db/db mice display not only deficits in the Morris water maze, but also impairments in long-term potentiation (LTP) and long-term depression (LTD) in the hippocampal area CA1 and administration of leptin does not ameliorate the deficits seen on the electrophysiological level (33). Moreover, when they grow up, the leptin receptor deficient db/db mice develop a permanent form of diabetes mellitus, whereas the leptin deficient ob/ob mice do not (23, 24) Interestingly, it was shown by using adult db/db mice, that diabetes impairs hippocampus-dependent memory, synaptic plasticity and adult neurogenesis, and that these changes in hippocampal plasticity and function are reversed when normal physiological levels of corticosterone are maintained, suggesting that cognitive impairment in diabetes may result from glucocorticoid-mediated deficits (34). This suggests that diabetes is a strong risk for the normal functioning and maintenance of neuronal processes.

It has been reported that obesity has an impact on human brain functions, e.g. by impairing performance when executive functions were tested (35, 36). Several studies have linked obesity to morphological alterations within the human brain. For example a linear association between higher BMI and smaller brain volume as well as reduced hippocampal volume has been shown (for details see (37)). Comparable to the situation in humans, the ob/ob mice display a reduction of ∼ 10 % in total brain volumes (Fig. 2). A comparable difference has also been noted by analyzing total brain weight of the obese mice in comparison to their controls (38, 39).

Among others, changes in adult neurogenesis, as well as in the morphology of individual neurons might contribute to obesity-induced brain volume reductions. Interestingly, the ob/ob mice showed deficits in adult hippocampal neurogenesis. The ob/ob mice display significant reduced numbers of PH3 as well as of DCX positive cells, indicating that not only cell proliferation is affected, but also differentiation into new neurons as well. Therefore, it was suggested that obesity may have an impact on adult neurogenesis. Indeed, rodents (mice (40) as well as rats (41)) that become obese due to a high-fat diet (HFD) also display impairments in hippocampal neurogenesis. Furthermore, in diabetic mice (db/db mice) hippocampal neurogenesis is not only impaired but dramatically reduced by up to 50 % (42). Thus, obesity may represent a risk factor for a reduction in adult hippocampal neurogenesis.

In addition to changes in adult neurogenesis, alterations in dendritic spines represent another morphological correlate of changes in neuronal plasticity (43). We noted slightly elevated (though non-significant) spine densities in the ob/ob mice in apical as well as basal dendrites of CA1 pyramidal neurons. Somewhat comparable to this finding, in a HFD mouse model, a significant increase in spine densities was noted (44). In contrast, diabetic db/db mice, display a strong reduction in dendritic spine densities in the dentate gyrus (42) as well as area CA1 (45), indicating that obesity in combination with diabetes results in a different neuromorphological phenotype.

Since the db/db mice did not only display striking morphological differences as compared to controls, it is not surprising that they also showed differences in their behavior as e.g. visible in the Morris water maze test (33). Based on this, we were interested in analyzing whether adult ob/ob mice also display altered behavior. In the open field the ob/ob mice covered less distance in the open field arena as compared to their lean controls. Likewise, db/db mice traveled less than age-matched controls (46).

The reduction in the movement as well as the fact that the ob/ob mice show less time rearing could be interpreted as anxiety-like behavior. Other studies that have used methods that allow to investigate anxiety-related behavior in more detail, could show that the ob/ob mice indeed display increased anxiety behavior. For example, the dark-light box as well as the elevated plus maze are considered valid tests for increased anxiety behavior in mice, because the animals naturally avoid open and bright spaces (47, 48). Ob/ob mice spent significantly less time in the illuminated area of the test arena in the dark light box compared to control mice, indicating increased anxiety like behavior (49). Ob/ob mice that were treated with repeated intraperitoneal injections of leptin and untreated ob/ob mice differ in the elevated plus maze performance. The leptin treatment was found to have ameliorated the anxiety related behavior (accompanied by a weight loss) in ob/ob mice (50). Thus, it is possible that leptin has anxiolytic capacities and that leptin-deficiency might contribute to anxiety related behavior. However, one should be cautious to interpret a lack of mobility of the adult ob/ob mice in the open field as anxiety-like behavior, since ob/ob mice already show reduced mobility even in their home cage (51, 52). Moreover, the validity of the open field as a readout for anxiety like behavior is discussed controversially (53). The ob/ob mice not only traveled less than control animals in the open field, they also display a reduced velocity. Thus, it is more likely that the adult ob/ob mice traveled less, due to their increased body weight and a comparable behavior is seen in humans (54).

In humans, an association between obesity and lower cognitive abilities was suggested to exist but this aspect is a matter of controversy. While in some studies impaired spatial learning of obese humans was reported (55, 56), in another study even better performance of obese humans in spatial learning tasks has been observed (57). Experimental evidence, however, strongly suggested an association of diabetes with cognitive decline and dementia (58). Similarly, in the MWM (Fig. 6-8) the obese ob/ob mice differ in their behavior from db/db mice that were found to display impaired spatial learning (33). The standard parameters evaluated in the MWM, such as distance or time to target, are highly dependent on mobility, which is e.g. reflected by swimming speed. The average swimming speed of ob/ob mice was significantly reduced (Fig. 6A), somewhat comparable to the velocity recorded in the OF (Fig. 5). Any behavioral analysis of animals aimed at testing cognitive function, always requires a similar capacity of mobility in order to achieve comparable results or a measurement of a parameter that is not biased by mobility. The parameter “swimming direction” is independent of the speed of this movement and allows monitoring movement towards the platform position. The analysis of this parameter clearly demonstrates a precise course towards the platform position for both groups of mice (Fig. 6C), indicating that the obese mice display no deficits in spatial orientation. Thus, at least in the present mouse model of obesity, and comparable to data obtained in humans (57), obesity does not seem to interfere with hippocampal-dependent spatial learning. To test the obese mice in an even more demanding setting of the MWM, the animals had to perform in a spatial reversal task, where the platform was moved to the opposite quadrant. This requires that the mice have to extinguish the previously learned goal position and acquire a path to the new position (30). The results for the spatial reversal task of the MWM in ob/ob mice corroborate the above described findings. Moreover, the obese mice show an even more accurate movement towards the platform position (Fig. 8C) combined with the reduced swimming speed (Fig. 8B). These results further give support for the notion that obesity does not seem to interfere with hippocampal-dependent spatial learning.

Addressing the causality of obesity and cognition in humans, previous studies yielded largely different results (21, 22). Among others, an inverse relationship between obesity and intellectual abilities has been postulated. Along this line of argument, a correlation between academic success and obesity has been reported (18, 19). However, by controlling the study for the social and economic background of the participants, no correlation between academic success and obesity was evident (20).

However, it is challenging to interpret and compare different studies properly, since methodological approaches as well as parameters controlled for (e.g. age, sex, education, comorbidities) differ between these studies (for more details see (21)). Animal models of obesity allow to control several parameters. Moreover, the models also allow to distinguish between the effects of obesity and obesity in combination with (developing) diabetes. The results obtained by using ob/ob mice, suggest that obesity does not have any impact on morphological alterations related to neuronal plasticity in the hippocampus and related hippocampus dependent behavior. However, we should keep in mind that obesity is a risk factor for diabetes and that there is strong evidence for an association of diabetes with cognitive decline and dementia in humans (58).

## Acknowledgement

We wish to thank Mrs Hanisch and Mrs Kaiser for excellent technical assistance. The study was supported by a grant of the European Regional Development Fund (ERDF; UHGWM 21).

## Statement on the welfare of animals

All applicable guidelines for the care and use of animals were followed.

